# Biophysical principles predict fitness of SARS-CoV-2 variants

**DOI:** 10.1101/2023.07.23.549087

**Authors:** Dianzhuo Wang, Marian Huot, Vaibhav Mohanty, Eugene I. Shakhnovich

## Abstract

SARS-CoV-2 employs its spike protein’s receptor binding domain (RBD) to enter host cells. The RBD is constantly subjected to immune responses, while requiring efficient binding to host cell receptors for successful infection. However, our understanding of how RBD’s biophysical properties contribute to SARS-CoV-2’s epidemiological fitness remains largely incomplete. Through a comprehensive approach, comprising large-scale sequence analysis of SARS-CoV-2 variants and the discovery of a fitness function based on binding thermodynamics, we unravel the relationship between the biophysical properties of RBD variants and their contribution to viral fitness. We developed a biophysical model that uses statistical mechanics to map the molecular phenotype space, characterized by binding constants of RBD to ACE2, LY-CoV016, LY-CoV555, REGN10987, and S309, onto a epistatic fitness landscape. We validate our findings through experimentally measured and machine learning (ML) estimated binding affinities, coupled with infectivity data derived from population-level sequencing. Our analysis reveals that this model effectively predicts the fitness of novel RBD variants and can account for the epistatic interactions among mutations, including explaining the later reversal of Q493R. Our study sheds light on the impact of specific mutations on viral fitness and delivers a tool for predicting the future epidemiological trajectory of previously unseen or emerging low frequency variants. These insights offer not only greater understanding of viral evolution but also potentially aid in guiding public health decisions in the battle against COVID-19 and future pandemics.

**Significance Statement:** This research presents a biophysical model that maps the molecular properties of SARS-CoV-2’s receptor binding domain into an epistatic fitness landscape. By linking the binding affinities of the virus to its epidemic fitness, we offer a powerful tool for understanding and predicting the emergence and success of new viral variants. Our model, validated with real-world data and informed by theoretical insights, provides a foundation for interpreting the evolutionary trajectory of past pandemics and predicting those of the future. The adaptability of this biophysical model extends to the key proteins of other viruses as well, signifying its potential in guiding public health interventions, and advancing our understanding of viral evolution.

## 1. Introduction

Since its emergence, the SARS-CoV-2 virus has undergone continuous genetic changes, giving rise to variants with increased transmissibility such as Alpha, Delta, and the recent Omicron. Each has contributed to significant surges in global COVID-19 cases. These genetic alterations in the viral genome have a profound impact on the structure and function of viral proteins, causing consequential changes in viral fitness (defined as the capacity of the virus to infect). Variants of concern (VoCs), such as Omicron BA.1, possess specific mutations in the spike protein. They have been linked to enhanced transmissibility [1, 2], augmented binding to host cell receptors, and heightened resistance to antibody neutralization. [3, 4]. Understanding the relationship between these mutations and viral fitness requires investigating their influence on molecular properties of affected proteins. Key viral proteins, like the receptor binding domain (RBD) of the spike protein, play a critical role in facilitating viral entry into host cells by binding to angiotensin-converting enzyme 2 (ACE2) [5], a functional receptor on cell surfaces. Furthermore, RBD serves as primary targets for the most potent SARS-CoV-2-neutralizing antibodies [6] and subject to evolutionary pressure from human immune system. Therefore, mutations on the RBD have been shown to be highly correlated with increases of fitness.

On the experimental side, Starr et al. [7] systematically scanned through every amino acid substitution in the RBD of the spike protein to determine the mutation effect on RBD folding and ACE2 binding and showed a substantial number of mutations are well tolerated or could even enhance ACE2 binding. In more recent research, Moulana et al. [8, 9] conducted a thorough examination of the binding affinity across all combinations of the 15 RBD mutations found in the BA.1 variant of SARS-CoV-2 in comparison to the original Wuhan Hu-1 strain. This exploration covered a total of 32,768 genotypes and involved testing against four monoclonal antibodies (LY-CoV016, LY-CoV555, REGN10987, and S309) as well as the ACE2 receptor. Additionally, global initiatives that promote data sharing, such as the Global Initiative on Sharing All Influenza Data (GISAID) [10], provide us with the ability to derive viral fitness based on prevalence data.

Prior studies have derived a quantitative correlation between fitness and molecular properties. For instance, Cheron et al. [11] and Rotem et al. [12] devised a theoretical framework to assess fitness of RNA viruses and validated it using experimental and computational methods. Central to their premise is that fitness comes from the proportion of viral capsid proteins in folded state free of antibodies, with state-occupation probability determined from Boltzmann distribution. SpikePro, a computational model, uses the spike protein’s amino acid sequence and structure to estimate SARS-CoV-2 fitness. The model considers the stability of the spike protein, its binding affinity with ACE2, and the potential for immune evasion [13]. While it has shown effectiveness in identifying dominant viral strains, it is worth noting that this is an empirical model whose foundation is not grounded in biophysical principles. Furthermore, experimental verification for the model’s predictions has been scarce, which highlights the need for more rigorous, physics-based models.

The central aim of our study is thus to bridge the gap between viral fitness and biophysical properties of the RBD. We concentrate specifically on how emerging mutations affect both fitness and the binding energies of the RBD to ACE2 and antibodies. By doing so, we aim to develop a robust methodology based on statistical mechanics to forecast RBD fitness anchored in its molecular properties.

This study establishes a biophysical link between binding affinities and relative fitness for SARS-CoV-2 mutants. To that end, we have constructed a genotype-to-fitness mapping for the SARS-CoV-2 RBD under the constraints of successful cellular entry via ACE2 and influence of neutralizing antibodies. This mapping equips us with a predictive tool for assessing fitness of emerging SARS-CoV-2 variants.

## 2. Results

### 2.1. The model

Our RBD fitness function is based on thermodynamics of protein folding and binding, as described in Figure 1. We consider the contribution of fitness (infectivity) from RBD to the virus, denoted *F*, as proportional to the fraction of RBD that are folded and free from antibodies.

**Fig. 1.**
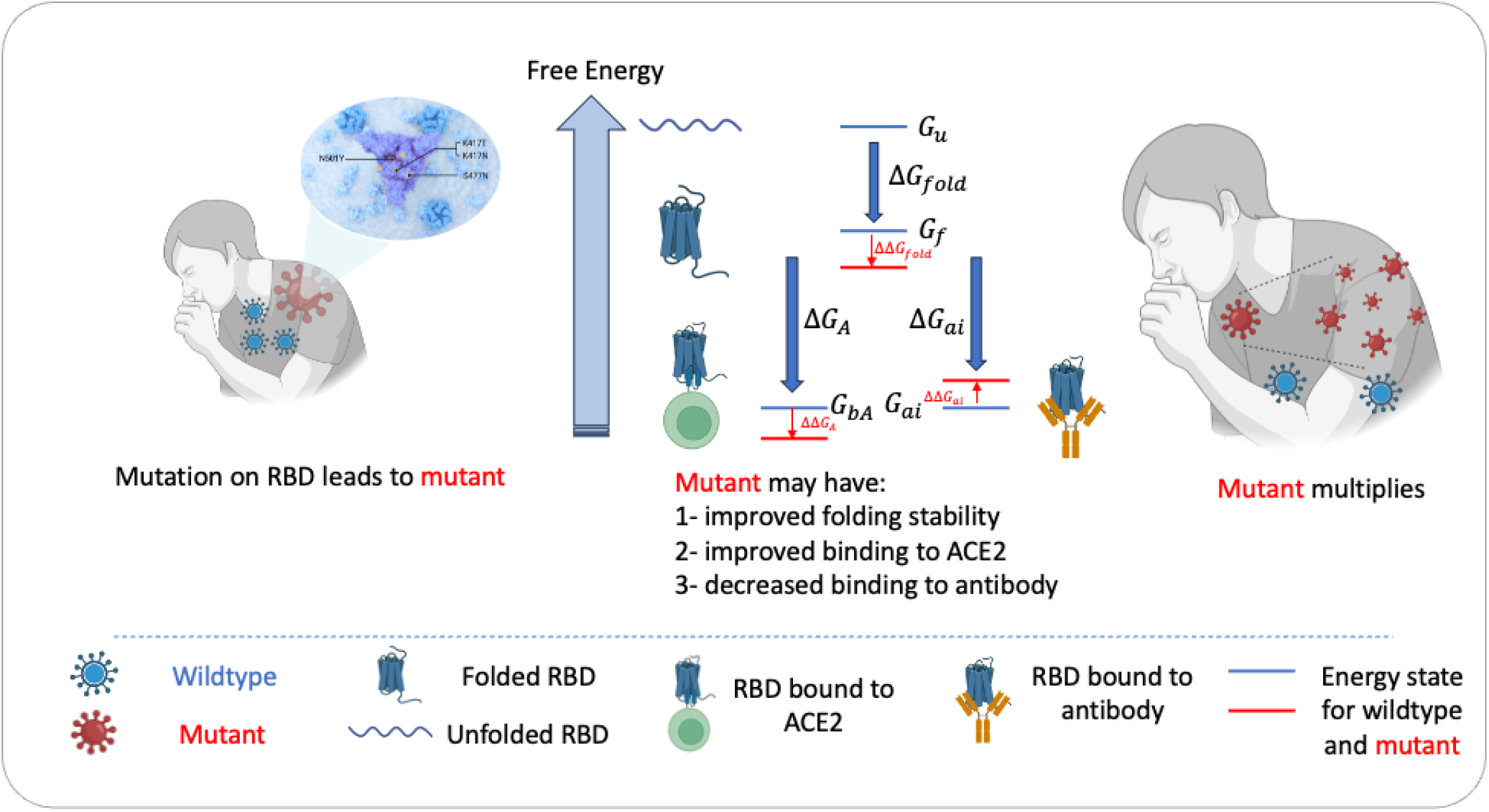
Model Illustration. Link between fitness and thermodynamics of protein folding and binding. High fitness variants may exhibit improved stability in the folded state or in the ACE2-bound state to facilitate cellular entry or have the capacity to destabilize bound-to-antibody states, thereby enabling evasion.

Specifically, we assume a seven-state microscopic configuration model for the RBD : unfolded, folded and unbound, folded and bound to ACE2, and folded and bound to one of four different antibodies, with respective free energies *G*_*u*_, *G* _*f*_, *G*_*bA*_ and *G* _*ai*_, where *i* is an index over the 4 antibodies. In our model, the RBD can only be bound to one antibody at a time, or ACE2 and to no antibody, or be free from ACE2 and antibodies. We then assume that an RBD can exist at thermodynamic equilibrium over these 7 states at some finite temperature *T*_*s*_, denoted by *β* = 1 /(*k* _*B*_*T*_*s*_), where *k* _*B*_ is the Boltzmann constant. Following [12], we then propose that fitness *F*, which in our context signifies the contribution to the viral infection rate from the RBD, is proportional to the Boltzmann probability of RBD being found in the folded state free from antibodies or in the folded, bound to ACE2 state. This approach is grounded in the understanding that while binding to the ACE2 receptor is crucial for initiating infection within an individual, the transmission of the virus between individuals is predominantly driven by ‘free viruses’ — those not yet bound to any receptors [14]. These free viruses, contained in the host’s body and airborne droplets, represent a potential state for initiating further infections in new hosts. Thus, fitness *F* can be expressed as

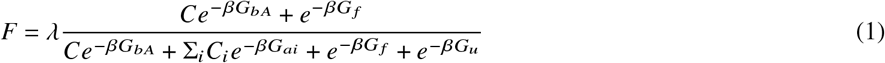

In this equation, 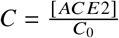 and 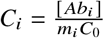 where […] represents concentration, *m*_*i*_ accounts for the quantity of antibodies required to neutralize the virus and *λ* is a scaling factor. The use of the standard reference concentration *C*_0_ allows us to express *C* and *C*_*i*_ in dimensionless form. For each mutant *mut*, the relative fitness compared to the wildtype *wt* (Wuhan-Hu-1), defined as 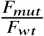, follows the same relationship as the absolute fitness. To avoid confusion, we will simply refer to this relative measure as the “fitness” *F*. Defining free energy differences between states as Δ*G* _*f old*_ = *G* _*f*_ − *G*_*u*_, Δ*G* _*A*_ = *G*_*bA*_ − *G* _*f*_ and Δ*G*_*ai*_ = *G* _*ai*_ − *G* _*f*_, and given that RBD domains are stable - with the unfolded states having significantly higher free energy than the folded states (that is, 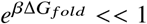, see “Methods”) - we can simplify the model as follows:

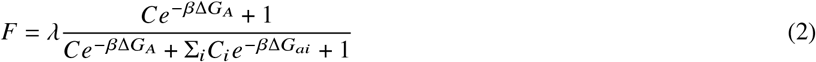

Given that Δ*G* ∝ *ln(K*_*D*_), we can express fitness as a logistic function of the logarithm of the dissociation constants *K*_*DA*_ and *K*_*Dai*_ for ACE2 and antibodies. Our study is centered on inferring *F* by fitting a scaling parameter *λ* and effective molecular concentrations in population *C* and *C*_*i*_ to the biophysical model. In our model, the temperature is treated as a hyperparameter that can be tuned (see “Methods” and SI for details). For clarity, a summary of all notations is provided in Table 1.

**Table 1.**
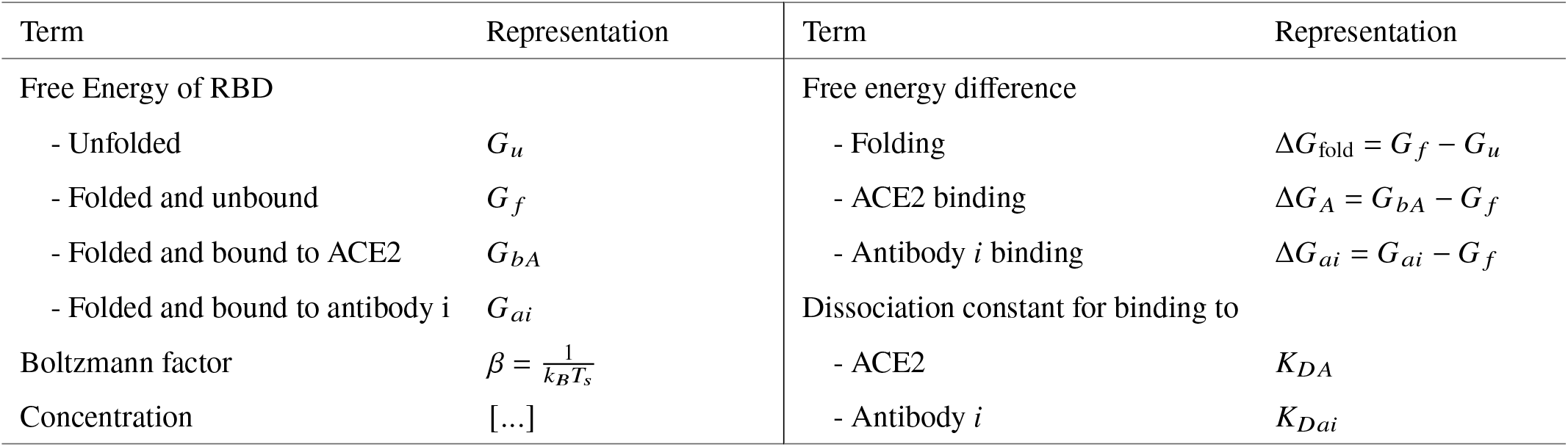
Terminology. A summary of terms and their representations used in the paper.

### 2.2. Fit of biophysical model to fitness obtained from population data

We separated our data into training and testing sets (see “Methods” for details) and then calibrated our biophysical model using observed viral variants [15] from the population study and incorporated experimental measurements of binding affinities [8, 9] as an input. Training of the model involved adjusting **6** parameters (*a, C*, and *C*_*i*_ for *i* ranging from 1 to 4) to achieve highest correlation between model prediction and fitness inferred from population data in the training set. During the fitting, Energy Scale T was fixed to 1.6, so that fitted concentrations (on the order ∼ 1 nM to ∼ 102 nM) agree with values of antibody concentrations observed in human serum (between 1 and ∼ 60 nM in severe symptoms) [16]. For the calculation of the effective concentration, we estimate that *m* is in the range of 10-100. [12] (see “Methods” and SI for details about fitting)

To demonstrate the predictive power of our biophysical model, we trained the model on 2% of the observed variants (22 points), achieving a highly satisfactory fit (average *R*^2^ = 0.97) on the training set as shown in Figure 2a. This result was further corroborated by the predictive power demonstrated on the test set with around 1000 variants (average *R*^2^ = 0.91), thus confirming the absence of over-fitting (Figure 2b). Our model’s performance reflects the biophysical understanding that a combination of strong ACE2 binding and reduced antibody binding from the four chosen antibody could provide the virus with a fitness strategic advantage, enhancing its ability to spread in the population.

**Fig. 2.**
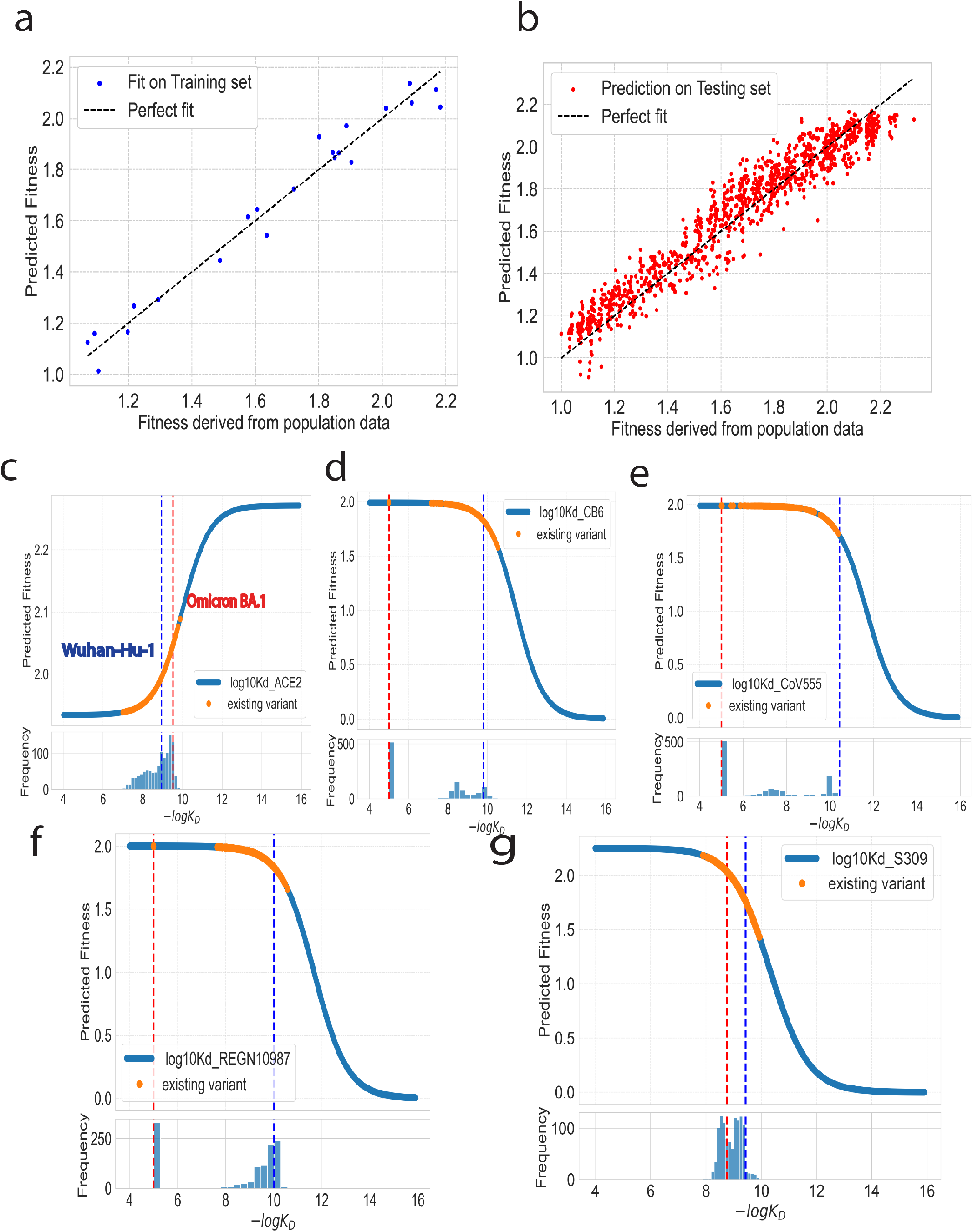
Biophysical Model Analysis. (a) Fit of the predictive model on the training set. Each dot represents a variant in the training dataset, plotted against the fitness derived from population data on the x-axis and the predicted fitness by the model on the y-axis (*R*^2^ = 0.97). (b) Model’s performance on the testing set. The model’s predictions align well with the fitness observed in population, as reflected by *R*^2^ = 0.94 suggesting that the model maintains a strong predictive power on unseen data. (c-g) The dependence of fitness function on the logarithm of each dissociation constant. Existing variants are depicted as yellow regions on the curve, and a histogram showing the distribution of the dissociation constants is provided beneath each plot. Red and blue dashed lines represent *wt* and BA.1 respectively.

To further explore the behavior of our model, we fixed all but one of the dissociation constants at their mean value across mutants and studied the variation of inferred relative fitness as a function of the unfixed constant. Interestingly, we observed that variants carrying a combination of Omicron mutations consistently fell into the upper plateau or linear segment of our biophysical model (Figure 2c-g). This suggests that natural selection did not favor combinations of mutations that would lead to high antibody binding. It should be noted that in each curve on Figure 2c-g, four out of five dissociation constants are fixed at the mean values. Thus, the inferred fitness for each data point in these one-dimensional cross sections of the fitness landscape does not represent its actual fitness.

### 2.3. Predicting RBD fitness for variants between wt to BA.1

Our model constitutes an effective tool for forecasting fitness contribution of RBD to the virus, given molecular properties of their RBD domain. We first examine the predictive power for variants between *wt* and BA.1.

In Figure 3a, the model is trained on the experimental dataset that excludes the G446S mutation and is subsequently tested on variants containing this specific mutation. Remarkably, our model even succeeds in predicting fitness of variants bearing the G446S mutation, a demanding task considering that this mutation causes escape from the REGN10987 antibody [9]. The model’s ability of predicting the impact of the G446S mutation on fitness, despite the complexity of correlating complete immune escape from REGN10987 to fitness, highlights the potential of our model to predict effects of unseen mutations.

**Fig. 3.**
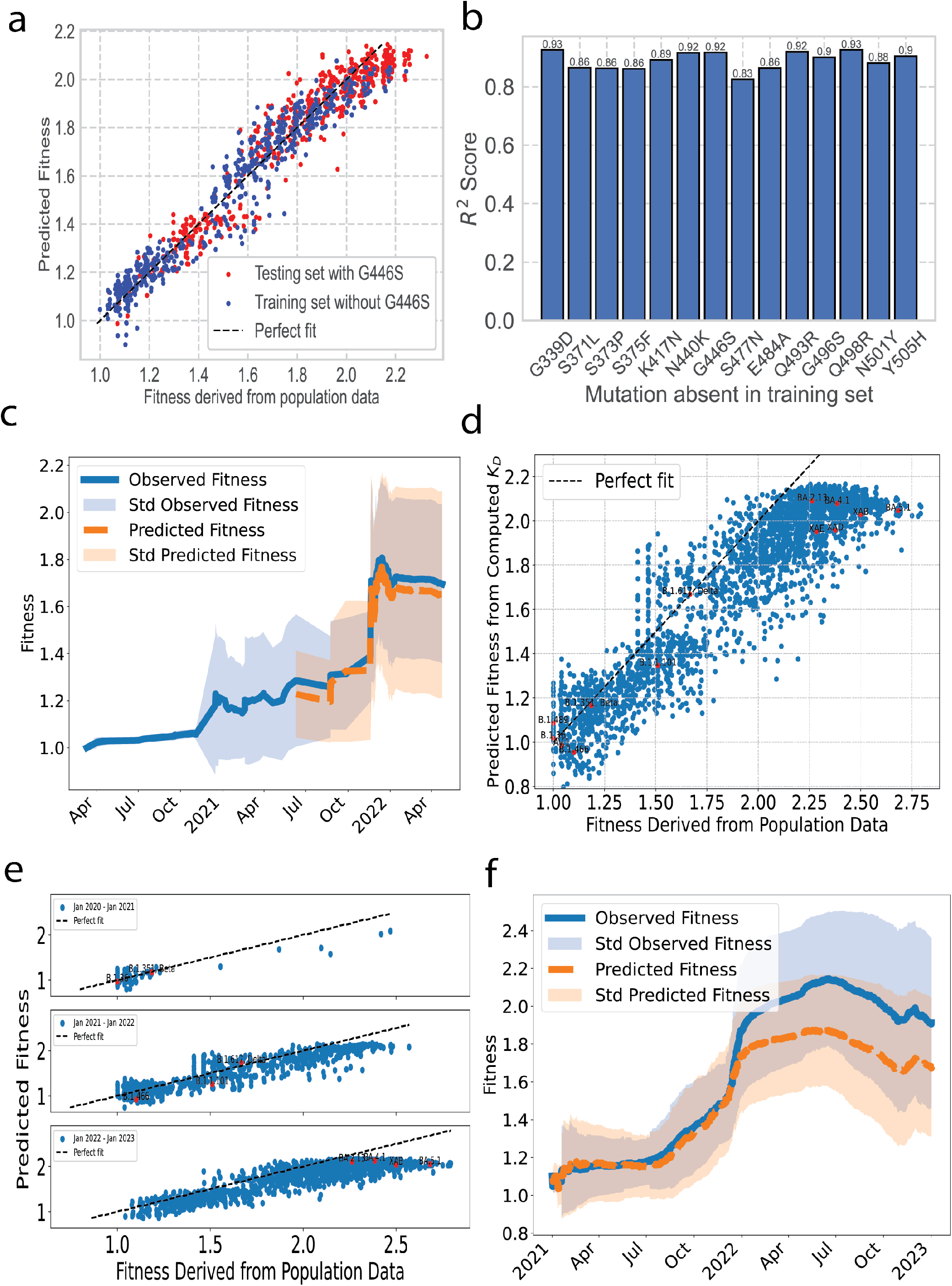
Assessing Predictive Power of the Model using experimental *K*_*D*_ (a-c) and computed *K*_*D*_ (d-f). (a) Fitness prediction for variants carrying the G446S mutation which was not included in the training set yields *R*^2^ = 0.92 (b) *R*^2^ derived from a model trained on variants excluding a specific mutation, then used to predict fitness of variants exhibiting that mutation. (c) Predictions of fitness compared with actual fitness trend for variants between Wuhan-Hu-1 and Omicron BA.1. Variants observed before May, 2021 are used as training set for the model. The model uses experimental *K*_*D*_ (d) Predicted fitness from biophysical model against actual fitness derived from population data. The model uses *K*_*D*_ acquired from ML and is fit on Wuhan-Omicron set. Selected variants are highlighted. (e) Predicted fitness compared with actual fitness for variants observed in 2020, 2021 and 2022. (f) Predictions of fitness over three-month rolling windows, compared with the actual fitness trends for variants between 2021 and 2023.

This point is further highlighted in Figure 3b. For each mutation, labeled as *m*, our model was systematically trained on all variants that exclude mutation *m* and then used to predict fitness of all existing variants carrying mutation *m* in combination with other possible mutations. This methodology, implemented across all 14 mutations, yielded a strong *R*^2^ coefficient of over 0.8. The efficacy of the model is demonstrated by its ability to consistently and accurately predict the fitness of previously unobserved mutation between Wuhan-Hu-1 and Omicron BA.1.

To further demonstrate the predictive capacity of our model, we assessed its ability to forecast fitness of future variants. The biophysical model was fitted on the dataset comprising all registered variants prior to May 01, 2021, which included only 42 data points. This trained model was then deployed to predict fitness of subsequent variants that emerged between May 1, 2021, and May 1, 2022, a period in which 843 unknown variants arose. Importantly, our model exhibited a notable predictive power, accurately predicting the leap in fitness induced by the Omicron BA.1 variant and its neighboring variants, as depicted in Figure 3c, with an *R*^2^ value of 0.76 (see SI). This underscores the model’s efficiency in predicting fitness of unseen variants, even when trained on relatively sparse data from the early stages of the pandemic.

### 2.4. Predicting RBD fitness beyond wt to BA.1

Moreover, while our model has been trained on mutations spanning from the Wuhan-Hu-1 to the Omicron BA.1, it’s not restricted to this specific spectrum. The model exhibits the capability to capture the effects of mutations outside the scope of Wuhan to Omicron, provided the relevant RBD’s biophysical properties are available, either through experimental data or simulations.

To estimate the fitness of variants beyond the experimental dataset of 215 sequences, it is essential first to determine the *K*_*D*_ values. We employed a supervised learning approach, utilizing transformer embedding and neural networks (see methods), proven effective in estimating the binding affinity of unseen RBD as demonstrated in Han et al. [17]. It is important to note that while the method we employed may not represent the most state-of-the-art *K*_*D*_ estimator available, its application here aims to illustrate how our biophysical fitness model can be synergistically integrated with either simulation or ML-based *K*_*D*_ estimators.

Our ML model is trained using 20,000 variants between Wuhan and Omicron BA.1 and their corresponding *K*_*D*_ values. The remaining 12,565 variants was then utilized for validation. On the validation set, we achieved *R*^2^ of 0.89, 0.98, 0.84, 0.75, and 0.79 respectively for ACE2 and four antibodies, as demonstrated in Figure S.1.

Then the ML model was applied to all RBD sequences observed in GISAID for which we could calculate population infectivity but lacked *K*_*D*_ values for the biophysical model. We estimate these *K*_*D*_ values using our ML model, then feed them into the biophysical model and compared against actual fitness metrics derived from population data as shown in Figure 3d. Importantly, this pipeline allows us to predict fitness for new variants, based solely on their sequences. Despite being trained on variants between Wuhan and Omicron BA.1, the model could correctly predict the fitness of variants with combinations of unseen mutations, achieving a Spearman correlation of 0.88. A selection of these variants are identified and labeled in Figure 3d for reference. Please note that for Figure 3d and subsequent panels e and f, variants between Wuhan-Hu-1 and Omicron BA.1, for which we already have experimental *K*_*D*_ data, are excluded from the analysis.

In Figure 3e, we assess the model’s capacity to predict RBD fitness for variants that emerged in 2020, 2021, and 2022. Our model not only accurately predicted the fitness of various variants but also consistently identified the top variants within each of these time frames. However, in 2023, a noticeable divergence emerged between the predicted and actual fitness, particularly for the Omicron sub-variants BA.2, BA.4, and BA.5. While our model successfully recognized them as the most infectious variants of the year, it was unable to discern the fitness differences among these sub-variants. We hypothesize that this limitation arises because the evolution of these variants are not predominantly driven by the four antibodies studied in this paper, given that BA.1 had already demonstrated escape from three out of the four antibodies. We then studied the model’s capacity to infer the fitness trend during the pandemic in Figure 3f. Although our model’s predictions start to diverge from the actual fitness metrics from early 2022 onwards due to the underestimation of the most fit variant, it maintains alignment with the general trend post-2022, showcasing its continued capability in accurately predicting fitness for other variants occurred during this period.

### 2.5. Epistasis

After fitting our biophysical model to all existing variants, we extrapolated fitness across all 32,768 possible mutation combinations which we have experimental *K*_*D*_ data. Notably, we observed a fitness threshold for the RBD, beyond which additional mutations cease to enhance fitness [18]. Essentially, once the virus achieves a certain fitness level, characterized by high immune escape and robust binding to the cell receptor, it becomes increasingly challenging for further mutations to improve this balance. As a result, fitness begins to plateau, as illustrated in Figure 4a. This phenomenon, which was not predicted by the work of Obermeyer et al. [15] or other non-epistatic models, can be attributed to two key factors. Firstly, as suggested by Moulana et al. [9], mutations that enhance antibody escape tend to reduce the virus’s affinity for ACE2 receptors. This indicates a trade-off in viral evolution between immune evasion and the ability to infect host cells, a factor inherently accounted for in *K*_*D*_ measurements and integrated into the biophysical model. Secondly, the logistic function used in our biophysical model naturally leads to a plateau in fitness.

**Fig. 4.**
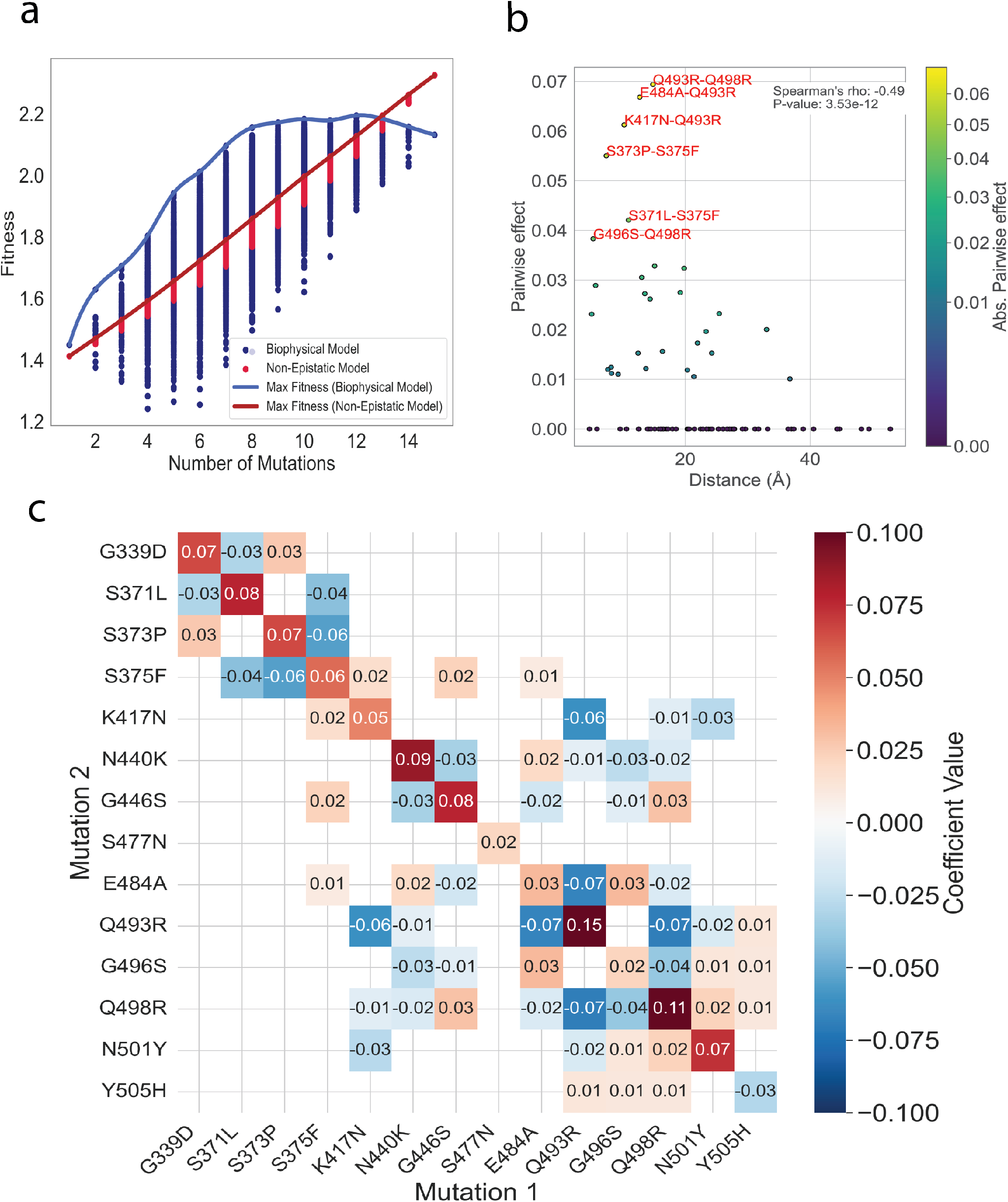
Epistasis Analysis. (a) Predicted fitness values with the non-epistatic model of Obermeyer et al. and with our epistatic biophysical model plotted against the genome’s mutation count, for all mutation combinations with mutation T478K. We define ‘Max Fitness’ as the maximum fitness prediction from our biophysical model. ‘Max Fitness’ curve begins to plateau with a higher mutation count, demonstrating a diminishing returns effect in epistatis. (b) Pairwise (second order) interaction coefficients against the spatial distances between the corresponding residues, with mutations colored in accordance with the absolute value of their pairwise coefficient. (c) Coefficients of epistasis: diagonal coefficients denote first order interactions, whereas off-diagonal coefficients represent second order interactions. Coefficients smaller than 0.01 have been masked for clarity.

To calculate the epistatic coefficients, we utilized a linear model described in Methods. We applied this model to fitness values inferred from our biophysical model across all 32,768 possible combinations of mutations. The performance of the model was evaluated (expressed as *R*^2^) for different orders of epistasis on a withheld test dataset, constituting 10% of the total data. A first-order model (comprising 16 coefficients) produced an *R*^2^ value of 0.958, while a second-order model (comprising 121 coefficients) achieved an *R*^2^ of 0.985. An F-test was conducted, yielding a p-value of 10*−*16. This confirms the robustness of the second-order model despite its increased parameter count over the first-order epistatic model. The high correlation values and satisfactory representation of the data suggest that a second-order epistatic model sufficiently captures the key dynamics, thereby alleviating the necessity for higher-order epistatic models. Following the training of the second-order model across the complete dataset to derive final coefficients, we observed that most single mutations (reflected on the diagonal of the matrix in Figure 4c) have a positive impact on fitness. An interesting aspect of our findings is the behavior of the Y505H mutation. While it displayed a negative first-order coefficient, its interactions with other mutations are positive, which effectively offsets the first order negative impact.

Furthermore, we observed a unique dynamic with the Q493R mutation. Alone, this mutation is beneficial, contributing to the virus’s escape from the LY-CoV555 antibody. However, its co-occurrence with other mutations reverses this benefit by reducing the binding affinity to ACE2 [19]. Particularly noteworthy are adjacent mutations in the crystal structure such as Q493R-E484A, Q493R-Q498R, and Q493R-K417N. These proximal mutation pairs demonstrate significant deleterious second-order effects, as illustrated in Figure 4b. Intriguingly, the evolutionary trajectory of Q493R seems responsive to these complex interactions. A reversal of the Q493R mutation was observed in subsequent lineages, including BA.4, BA.5, BA.2.75, BQ.1, and XBB. This reversal suggests an evolution dynamics driven by the trade-offs between immune escape and ACE2 binding.

Overall, these findings highlight the complex dynamics of viral evolution, where multiple mutations interact non-linearly to enhance or reduce viral fitness.

## 3. Discussion

Although the complexity of fitness landscapes is undeniable, recent research indicates that in certain biologically and clinically significant systems, such as evolution of bacterial resistance against antibiotic [20], evolution of viral resistance against antiviral treatments [11, 21], as well as norovirus evolution against a neutralizing antibody [12], these fitness landscapes can be systematically and quantitatively delineated. Our research builds upon these findings and reveals that the fitness landscape of the SARS-CoV-2 RBD, undergoing evolution against neutralizing antibodies, can be systematically described through its biophysical properties. In our study, we found the strength of binding to the neutralizing antibody, as well as to ACE2, play crucial roles in determining RBD fitness. Importantly, the significance of these biophysical parameters is not confined to SARS-CoV-2 alone. Similar traits, namely antibody binding affinity and protein folding stability, have been crucial in influencing the evolution of influenza viruses [21–23]. This observation suggests that our model may have broader applications, potentially extending to other viruses beyond SARS-CoV-2.

To forecast the fitness of emerging novel variants with our model, acquiring the dissociation constants for the corresponding mutated RBD is imperative. While experimental data sets the gold standard, obtaining them early in a pandemic may pose challenges. Fortunately, computational methodologies have demonstrated efficacy in predicting binding constants for unobserved mutations using molecular dynamics (MD) and ML. For instance, Lacam et al. [24] achieved exceptional accuracy by employing a framework that integrates MD and potential-of-mean-force calculations. Their study accurately determined the binding free energy of the RBD for four prevalent variants and the wild type, when complexed with ACE2 or antibodies S2E12 and H11-D4. In parallel, Sergeeva et al. [25] offered an effective strategy to determine the impact of interfacial mutations on the binding affinities between RBD and ACE2, using free energy perturbation. Williams et al. [26] constructed a multi-layer neural network, using biophysical parameters as inputs to predict binding affinities of SARS-CoV-2 antibodies with various VoCs. Similarly, Chen et al. developed the NN-MM-GBSA model [27] using MD and neural network to predict binding affinity between SARS-CoV-2 spike RBD variants and ACE2, reaching a correlation coefficient of 0.73 on prediction of dissociation for single variants in the work Starr et al. [7].

In this study, we estimated binding affinities using a computationally efficient alternative by employing a neural network that takes in transformer embedding for sequences and outputs *K*_*D*_. In this way, we could predict mutational effects not only for combinations of mutations outside of experimental data set but also for completely unseen mutations. By incorporating these predicted binding constants into our biophysical model, we demonstrate that this streamlined computational approach yields accurate predictions for VoCs both at the early and later stage of the pandemic.

During the early stages of a pandemic, data on viral fitness or infectivity is typically limited. Unlike *K*_*D*_ measurements, which can be derived from wet lab experiments or simulations, comprehensive fitness data is usually only accessible after a variant has extensively spread and been subjected to population-level sequencing. Therefore, the ability of our model to train and predict effectively with minimal data points is particularly valuable in the context of pandemic preparedness. Considering the RBD’s susceptibility to mutations, our model has the potential to be a powerful tool in understanding and predicting the fitness of emerging variants.

Despite our biophysical model being trained on non-epistatic population data from Obermeyer et al. [15], the model allows us to generate an epistatic map from genotype to fitness using *k*_*D*_. In our model, we observe a tendency for fitness to plateau in the face of increasing number of mutations relative to the wild type. This phenomenon of diminishing returns, manifested as a fitness plateau, has been extensively studied in existing literature [18, 28–31].

Our results also indicate that epistasis constrains evolution. While many of the mutations we investigated are beneficial, specific combinations of mutations could be deleterious. Some mutations require the concurrent occurrence of stabilizing mutations to counterbalance their adverse consequences. This observation is consistent with findings of Gong et al. [21] and Rodrigues et al. [20] underscoring the critical role of stabilizing mutations in fixation of subsequent destabilizing mutations that could hold adaptive value.

Our results emphasize the importance of accounting for the interactive effects of multiple mutations in viral evolution modeling and prediction. It is particularly noteworthy that our model could explain the reversal of Q493R from a viral fitness perspective. Furthermore, in our model, second-order pairwise effects between mutations tend to weaken as their separation in the crystal structure increases. These observations demonstrate that our model effectively captures the essential aspects of viral fitness, including the epistasic effects that drive viral evolution.

A fundamental assumption of our model is that evolution of the SARS-CoV-2 RBD is predominantly driven by its capacity to bind ACE2 receptor and evade antibodies. Our model, which is based on this assumption, demonstrates the ability to predict fitness effect of all mutations in RBD, except for T478K. T478K is an interesting mutation as shown by Moulana et al. [8, 9] that it had a negligible effect on dissociation constants, despite its strong contribution to variant infectivity. The exact reason for T478K’s high fitness contribution, despite no apparent change in RBD’s biophysical parameters, remains unclear. A prevailing hypothesis is its frequent co-occurrence with the D614G mutation [32]. D614G, located on the spike protein but outside of the RBD, has been shown to enhance viral replication and infectivity [33, 34]. Our model, focusing on the RBD, does not account for effects of mutations like D614G on the spike protein. This limitation leads to our observation that the RBD containing T478K consistently shows increased fitness, regardless of the mutational background in RBD. Consequently, we segregated the training data based on the presence of the T478K mutation. This approach helps us account for the fitness increase associated with T478K, despite our model’s focus on the RBD.

Research, including that by Lee et al. [35], has demonstrated that mutations on the RBD could alter the conformation of the Spike trimer, affecting the proportion of RBDs in the ‘up’ state, which in turn enhances infectivity by facilitating ACE2 binding. To adequately consider the ‘up’ and ‘down’ states of the RBD, it is crucial to acquire binding affinity measurements on the entire spike trimer, as demonstrated in the studies by Xu et al. [36] and Yin et al. [37] on specific VoCs, rather than on isolated RBD. *K*_*D*_ measurements on spike would capture the effective binding affinities averaged over the two configurations at equilibrium, leading to a similar formulation as Equation 1 but with binding affinities to the Spike protein instead of the isolated RBD. Our current model, however, does not incorporate these aspects as the *K*_*D*_ are measured using isolated RBD with yeast display. Nevertheless, studies like those by Costello et al. [38] have indicated that RBD dynamics in isolation are comparable to those in the intact spike, which forms the basis for numerous biochemical studies using isolated RBD.

On the other hand, extrapolating our fitness predictions beyond RBD to encompass the entire viral sequence, presents a more formidable challenge. Our model’s predictive power on RBD fitness comes from the fact that binding affinity to ACE2 and antibodies are major evolutionary forces that drive RBD’s evolution. It is likely that mutations outside the RBD will have functional and structural effects that extend beyond alterations to binding constants with cell receptors and antibodies, complicating the predictions.

Furthermore, our model currently considers antibody binding to ACE2 and four monoclonal antibodies:LY-CoV016, LY-CoV555, REGN10987, and S309, an oversimplification given the complexity of human immune responses. While our model already exhibits accurate predictions and could easily be extended to other antibodies if data is available, the inclusion of additional factors such as other antibodies, vaccination effects, the replicative capacity within the infected cell [39], transmission dynamics [40], and drug resistance [41] could potentially enhance its predictive power and realism.

## 4. Methods

### 4.1. RBD Fitness Data analysis

We acquired fitness ratio of each RBD compared to wild type from the work of Obermeyer et al. [15]. In their study, fitness label is obtained by modeling the relative growth rate of SARS-CoV-2 lineages using a hierarchical Bayesian regression model. The model combines individual mutations and clusters genetically similar genomes to estimate the incremental effect of amino acid changes on growth rate within each lineage, which enables the model to share statistical strength among similar lineages. Specifically, the proportion of lineages is modeled as a multinomial distribution whose probability parameter is a multivariate logistic growth function softmax (*α* + *tb*/*τ*). For each lineage, the slopes *b* are linearly regressed against the presence of each possible amino acid substitution *X*_*m*_ ∈ {*0*, 1} as *b* =Σ_*m*_ *b*_*m*_*X*_*m*_. These linear coefficients *b*_*m*_ can be directly interpreted as the effect of a mutation *m* on a lineage’s fitness. This model assumes each single point mutation independently linearly contributes to change in fitness. Authors reported that fitting a similar model of both single and pair mutations lead to no pairwise mutations stronger than the top 100 single mutations.

This enabled us to estimate fitness *F*_*mut*_ of each existing RBD mutant, compared to wildtype. Using 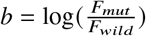, we get:

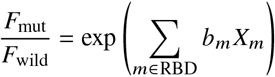

### 4.2. RBD Binding Affinity

We acquired the binding affinity data from the work of Moulana et al. [8, 9] In their study, they systematically examined the interactions between all possible combinations of 15 RBD mutations (totaling 32,768 genotypes) and ACE2, as well as four monoclonal antibodies (LY-CoV016, LY-CoV555, REGN10987, and S309) via Tite-seq measurement. In situations where binding affinities in their dataset were too low to measure accurately, we have chosen to substitute these with a fixed value of 5. We stress that this choice of value is not expected to influence our study’s outcomes. As indicated in Figure 2c-g, the antibody escape largely resides in the upper plateau region of fitness curve, thus, this preset value for variants with immune escape to antibodies does not have a substantial impact on the logistic regression results. Furthermore, we eliminated approximately 100 genotypes from the analysis that did not have measured ACE2 binding affinity.

### 4.3. Effect of mutations on RBD stability

In the presented results, we excluded the unfolded state from the model, on the assumption that this variable shows minimal variation across different variants and hence, would not significantly influence fitness. To verify this assumption, we employed DDGUN, an untrained, high-throughput tool [42], to compute ΔΔ*G* _*f old*_, the variance in folding free energy difference, for mutants relative to the wild-type: the maximum variation was under 2*k cal mol*. Recalling that Δ*G*_*f old*_ ≈ − 10 kcal/mol [43], we deduce that most mutations do not significantly destabilize RBD.

This could be indicative of the universally efficient folding of RBDs seen in nature. The selection pressure acting on the RBD primarily focuses on binding to ACE2 and immune evasion, and therefore the mutations are predominantly on the protein surface and do not significantly affect the protein’s stability.

### 4.4. Filtering RBD sequences from GISAID

All 15,371,428 spike sequences on GISAID [10] as of 14-April-2023 were downloaded and aligned, following approach in the work of Starr et al. [44]. Sequences from non-human origins and with lengths outside [1260, 1276] were removed. They were then aligned via mafft [45] and sequences containing unicode errors, gap or ambiguous characters were removed. Overall, we retained 11,976,984 submissions represented by 25,725 unique RBD sequences. RBD amino-acid mutations were enumerated compared to the reference Wuhan-Hu-1 SARS-CoV-2 RBD sequence (Genbank MN908947, residues N331-T531).

We then remove all RBD sequences that do not match any of the possible intermediates between Wuhan Hu-1 and Omicron BA.1. To do this, we allow all possible combination of 15 mutations (G339D, S371L, S373P, S375F, K417N, N440K, G446S, S477N, T478K, E484A, Q493R, G496S, Q498R, N501Y, Y505H) between Wuhan and Omicron BA.1, which lead to 215 = 32, 768 possible combinations. We calculated the number of occurrences of each RBD sequence as well as the time of its first occurrence, which we approximated by taking 5 % quantile of time data for each RBD sequence. From this analysis, we obtained 1121 unique observed RBDs. These RBDs correspond to 2.5 million sequences out of the 12 million sequences we initially screened from the GISAID database.

### 4.5. Fitting the model with logistic regression

For the purpose of logistic regression analysis, we utilized the intersecting data obtained from “Methods” RBD Fitness and RBD Binding Affinity. This accumulated dataset comprises 1118 unique RBDs observed in the GISAID database [46], for which the *K*_*D*_ values have been experimentally determined. We further partitioned this data based on the presence or absence of the T478K mutation within the sequence, resulting in two distinct subsets. (see Supplementary Information and Discussion).

The ratio between system temperature *T*_*s*_ (corresponding to body temperature of the host) and experimental temperature *T*_*e*_ (corresponding to the temperature of experiments conducted in work from Moulana et al. [8, 9], leading to Δ*G* = *RT*_*e*_*in* (*K*_*D*_)) was treated as a hyperparameter T (simply referred as “Energy Scale”), which is a parameter whose value is chosen before the fit is done.

We calculated unknown parameters *a, C* and *C*_*i*_ by fitting:

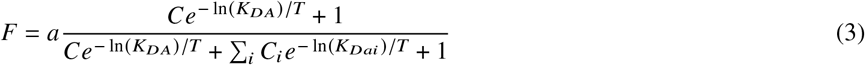

where Energy Scale T is fixed to 1.6. We emphasize that the hyperparameter can be chosen with knowledge of the training set alone (and thus does not invalidate prediction capabilities of the model) and that the model behavior is only slightly affected by changes in this hyperparameter (see SI for model performance at different Energy Scales).

Model fitting was performed with non-linear least square regression (*scipy*.*optimize* package) on a randomly selected training set. The model is then evaluated on the remaining testing set to prove absence of overfitting. Additionally, to mitigate the effects of randomness, we implemented 10-fold cross-validation wherever feasible.

### 4.6. Fitness prediction using ML estimated K_D_

Considering the potential unavailability of experimental measures for dissociation constants early in a pandemic, we advocate for the application of our methodology to *K*_*d*_ estimates derived from a computationally inexpensive deep learning pipeline. Notably, Hie et al. successfully predicted viral escape using a machine learning technique designed for natural language processing [47]. Similarly, Han et al. introduced an online platform based on deep learning models, incorporating transformers, for rapid prediction of binding affinity between RBD mutants and ACE2 [48].

Building on these developments, our study showcases a framework that combines a biophysical model with a *K*_*D*_ predictor utilizing protein sequence embedding and a neural network. This integrated system efficiently predicts dissociation constants for emerging variants, and the results are directly fed into the biophysical model. This method enables prompt and effective prediction of viral evolution in response to novel mutations.

ESM-1v [49] is a Transformer-based language model specifically trained on a diverse dataset of 98M protein sequences. This pre-trained model inputs a given protein sequence and outputs a vector representation, or an ‘embedding’, of that sequence. This embedding consists the evolutionary information of the protein sequence, which is pivotal in enabling the machine learning model to predict the effects of various combinations of unseen mutations as demonstrated in these papers [17, 47, 49].

To elaborate the embedding mathematically, consider a protein sequence of length *L*, we describe it as a sequence of tokens 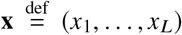. For the RBD, we have *L* = 201. During the forward pass of ESM-1v, we obtain the hidden representations from the final layer, denoted as (**h**_1_, …, **h**_*L*_), with each **h**_*i*_ being a vector in ℝ^1080^. To generate a comprehensive representation of the entire sequence, we apply mean pooling to these vectors, resulting in a single sequence representation 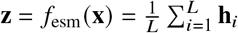.

After acquiring the embedding, we trained a neural network that takes the embedding as input and output the variant binding affinity with ACE2 and four antibodies. This network was trained using *K*_*D*_ data from 20,000 variants and subsequently validated on a separate set comprising the remaining 12,565 variants.

After training, the neural network was deployed to estimate the *K*_*D*_ for existing variants with known population fitness from section 4.1, encompassing a total of 5,460 variants. Excluding the variants between Wuhan and Omicron BA.1 (for which we already have experimental binding affinities), we focused on evaluating the predictive power of the remaining 3,334 variants. These estimated *K*_*D*_ values were subsequently integrated into the biophysical model, as described in section 4.5.

### 4.7. Epistasis analysis

Epistasis describes how mutation interactions can affect fitness *F*. If there is no epistasis then fitness can be described as a linear combinations of the presence of each mutation *X*_*m*_ ∈ {*0*, 1}, leading to a first order epistatic model:

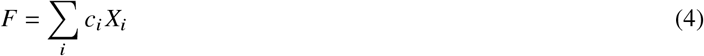

If we consider pairwise epistatic interactions between mutated sites, we get second-order epistatic model:

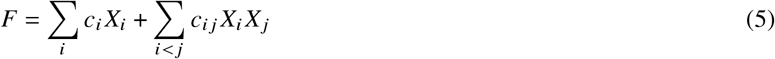

*c*_*i*_ are considered as “first-order” epistatic coefficient while *c*_*i, j*_ are “second-order” epistatic coefficients as they illustrate the non-linear epistatic interaction between mutated sites *i* and *j*.

To make sure the linear model does not overfit and can generalize on unseen data, we implemented a 10-fold cross-validation strategy (dataset split: 90% / 10%) and identified a linear model involving first and second order coefficients as described in Equation 5 gives a better representation of data than a first order model (*R*^2^ = 0.98 vs *R*^2^ = 0.96 on test set).

To analyze relationship between second order epistatic coefficients *c*_*i, j*_ and distance between mutated sites, we calculated the latter as the Euclidean distance between the average position (computed as the mean of positions of all non-Hydrogen atoms in the amino acid) of each mutated site.

## Acknowledgements

This work is supported by NIH R35GM139571 (to E.I.S) and NIGMS T32GM144273 (to V.M.). The content is solely the responsibility of the authors and does not necessarily represent the official views of the National Institute of General Medical Sciences or National Institutes of Health.

We gratefully acknowledge all data contributors, i.e., the Authors and their Originating laboratories responsible for obtaining the specimens, and their Submitting laboratories for generating the genetic sequence and metadata and sharing via the GISAID Initiative, on which this research is based. We would like to thank Vaibhav Upadhyay, Krishna Mallela, Zechen Zhang and Junlang Liu for useful discussions.

## Author contributions

D.W., M.H., V.M., and E.I.S. conceived and designed the research approach. D.W., M.H., and V.M. executed the analysis. E.I.S. supervised the work. All authors, D.W., M.H., V.M., and E.I.S., contributed to writing and reviewing the manuscript.

## Data and Materials Availability

All code used for the analyses in this study can be accessed on GitHub at https://github.com/Dianzhuo-Wang/RBD_fitness/. Any additional data or materials related to this paper can be obtained upon request from the authors.

