## Supplementary Material for "Biophysical principles predict fitness of SARS-CoV-2 variants"

### Supplementary Materials for:
### Biophysical principles predict fitness of SARS-CoV-2 variants

Dianzhuo Wang et al.

#### Supplementary information

This file includes:

Section S1. Machine Learning Prediction of $K_D$ on validation set
Section S2. Linear approximation and contributions to fitness
Section S3. Robustness of the model
Section S4. Energy Scale as a hyperparameter
Section S5. Prediction of future variants
Section S6. Justification of the split of the data using T478K and biophysical implications
Fig. S1. ML-predicted $K_D$ against experimental value s
Fig. S2. Linear approximation for logistic model
Fig. S3. Contributions of mutations to fitness
Fig. S4. Fitted parameters of the model
Fig. S5. Effect of Energy Scale hyperparameter
Fig. S6. Prediction of future variants between Wuhan-Hu-1 and Omicron BA.1
Fig. S7. Comparison of predicted fitness vs true fitness for existing variants with and without Mutation T478K

#### *S1. Machine Learning Prediction of $K_d$ on validation set*

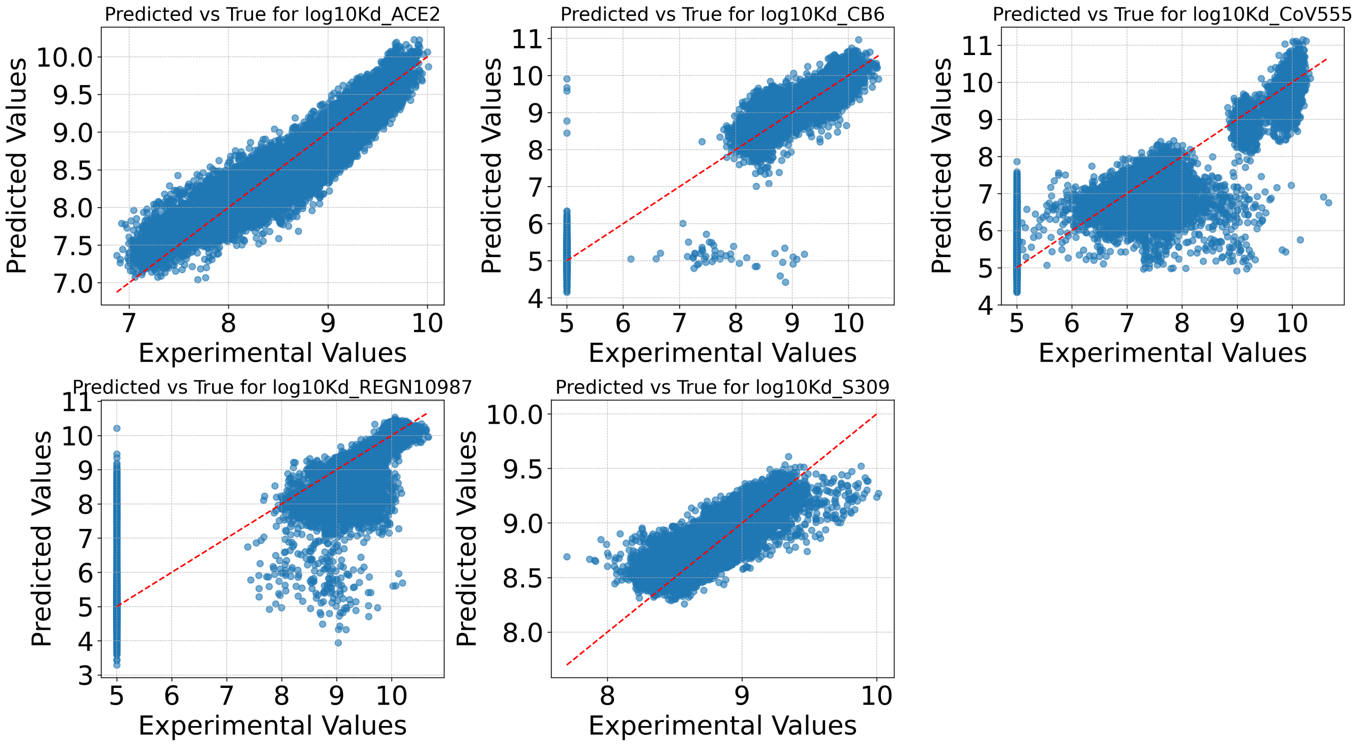


Fig. S.1. Comparison of ML-predicted $K_D$ values against experimental $K_D$ values for the validation set. The training set comprises experimental $K_D$ values for 20,000 variants, while the validation set includes the remaining 12,565 variants. The $R^2$ values for each of the features are as follows: 0.89, 0.98, 0.84, 0.75, and 0.79, respectively.

#### S2. Linearization and contributions to fitness

To convert improvement of binding to ACE2/immune escape (i.e. change in binding constants) of variants compared to wildtype into a fitness change, we fit a linear regression $\hat{F} = \alpha_0 + \sum_{i\in\{A,ai\}} \alpha_i log(K_{Di})$ to the observed variants.

This approach can be viewed as a linear approximation of a hyperdimensional sigmoid function, providing a simplified representation of the fitness landscape. A one-dimensional cross-section of this approximation is illustrated in Fig. S.2

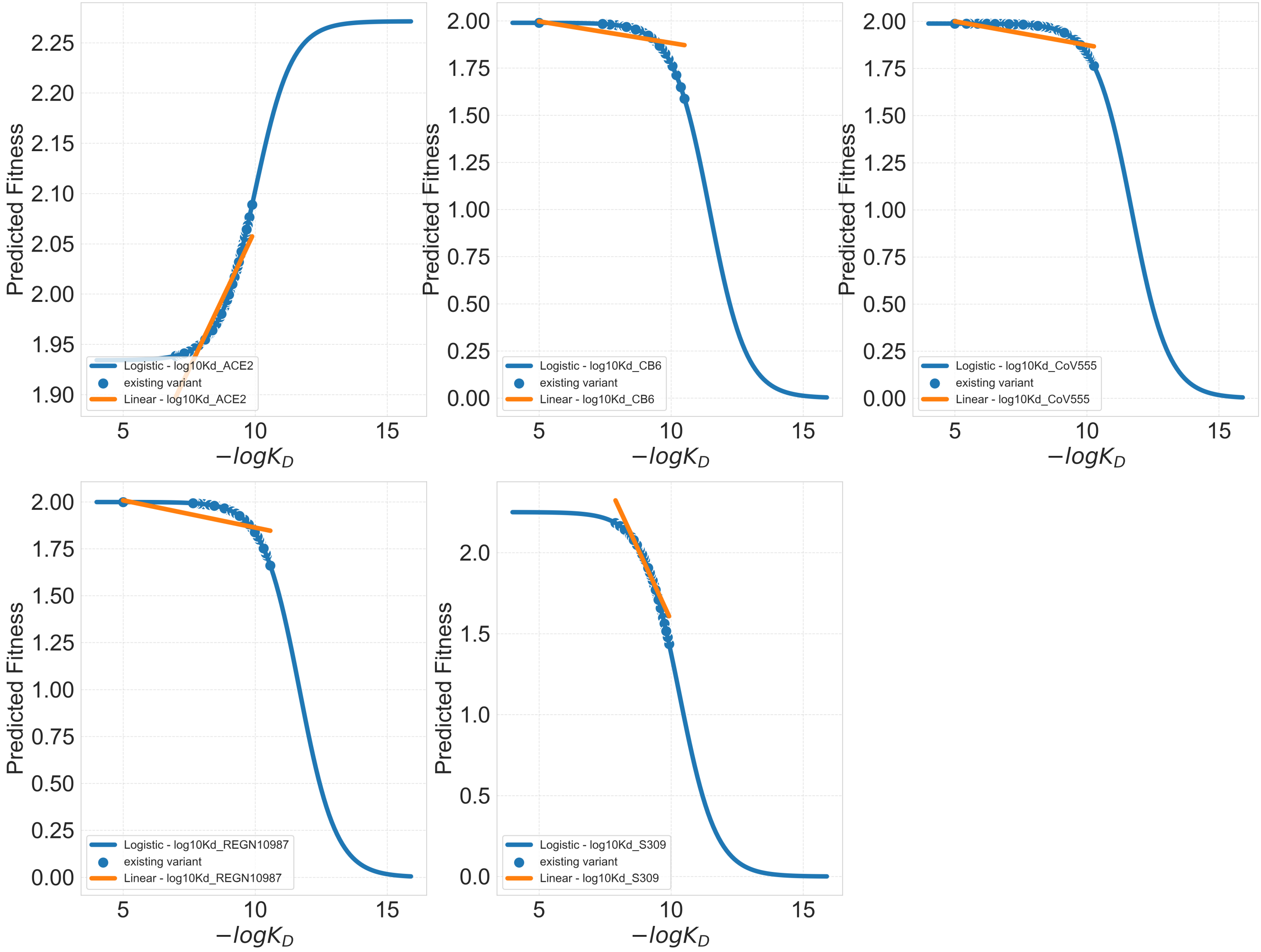


Fig. S.2. One-dimensional cross-sections of the logistic model alongside their linear approximations are presented. In each subfigure, four out of five dissociation constants are held constant.

To show how scaling of biophysical constant leads to fitter variants, we first computed the difference in biophysical constant, for each variant, compared to wildtype (eg.$\Delta \log(K_{Di}) = \log(K_{Di,mut}) - \log(K_{Di,wild})$), and then multiplied this quantity by the corresponding coefficient $\alpha$ in linear regression. The result for each variant is interpreted as a fitness increase (eg. $\Delta f_{\log(K_{Di})} = \alpha_{\log(K_{Di})}.\Delta \log(K_{Di})$). We then estimate the contribution of each mutation to the relative fitness increase by regressing it against substitutions features. Formally, we fit for each biophysical constant $\log(K_{Di})$: $\Delta f_{\log(K_{Di})} = \sum_{m\leq 15} \epsilon_{m,\log(K_{Di})} X_m$ with $X_m \in \{0,1\}$ where $\epsilon_{m,\log(K_{Di})}$ can be interpreted as the contribution of biophysical constant $\log(K_{Di})$ to fitness increase given by mutation $m$. Fitness difference between variant and wildtype is $\Delta F = \sum_{\log(K_{Di})} \Delta f_{\log(K_{Di})} = \sum_{\log(K_{Di})} \sum_m \epsilon_{m,\log(K_{Di})} X_m = \sum_m (\sum_{\log(K_{Di})} \epsilon_{m,\log(K_{Di})}) X_m$. Thus, $\sum_{\log(K_{Di})} \epsilon_{m,\log(K_{Di})}$ represents the contribution of mutation $m$ to fitness and $\epsilon_{m,\log(K_{Di})}$ the projection of this contribution on each feature.

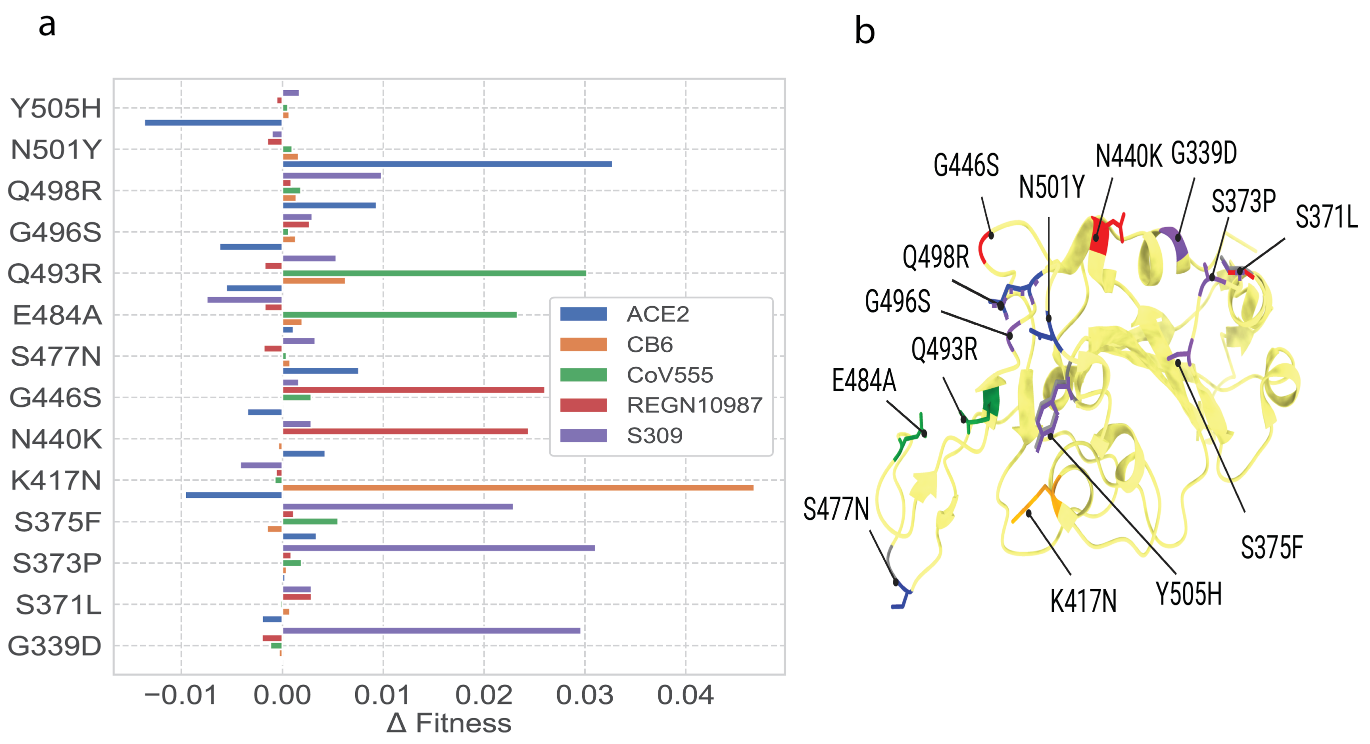


Fig. S.3. (a) Contributions of each mutation to fitness either towards evading specific antibodies or facilitating cellular entry. (b) A crystal structure of the wild-type RBD, with Omicron substitutions indicated. Mutated residues are color-coded based on the largest fitness contribution associated with the mutation. Bi-color representations suggest that two features contribute equally to fitness.

Through the linear approximation of our biophysical model, we can translate the observed improvements in ACE2 binding or immune escape (i.e., changes in binding constants) of specific variants compared to the wild type into alterations in fitness effect for each individual mutation. In Figure S.3 (a), we have quantified the individual contributions of each mutation towards either the evasion of specific antibodies or the binding to ACE2, thus unravelling the complex landscape of viral adaptation strategies.

In Figure S.3 (a), we observe that fitness effects for each mutation is primarily associated with one or two biophysical parameters. Furthermore, mutations spatially close on the spike protein often share similar characteristics contributing to their fitness. This observation is graphically represented on the crystal structure in Figure S.3 (b).

##### S3. *Robustness of the model*

Validation of the robustness of our model was undertaken by applying it to ten non-overlapping subsets of observed variants (Figure S.4). The error bars illustrate minor variations in coefficients, indicating the model's stability. Moreover, the coefficients for datasets, both with and without the T478K mutation, were remarkably similar. Intriguingly, the effective concentrations derived from our model for antibody binding demonstrated remarkable consistency across LY-CoV016, LY-CoV555, and REGN10987. A notable exception was the coefficient corresponding to the S309 antibody, which continues to be effective across a range of variants, from the original Wuhan strain to Omicron BA.1, thereby preventing viral immune escape. This unique attribute of S309 is represented in our model as a relatively higher effective concentration. To obtain the logistic regression coefficients, we fitted our model on the whole dataset of existing variants (for which we have fitness coming from population data). We computed error bars as the standard deviation of coefficients of the model fitted on 10 non-overlapping subsets of existing variants.

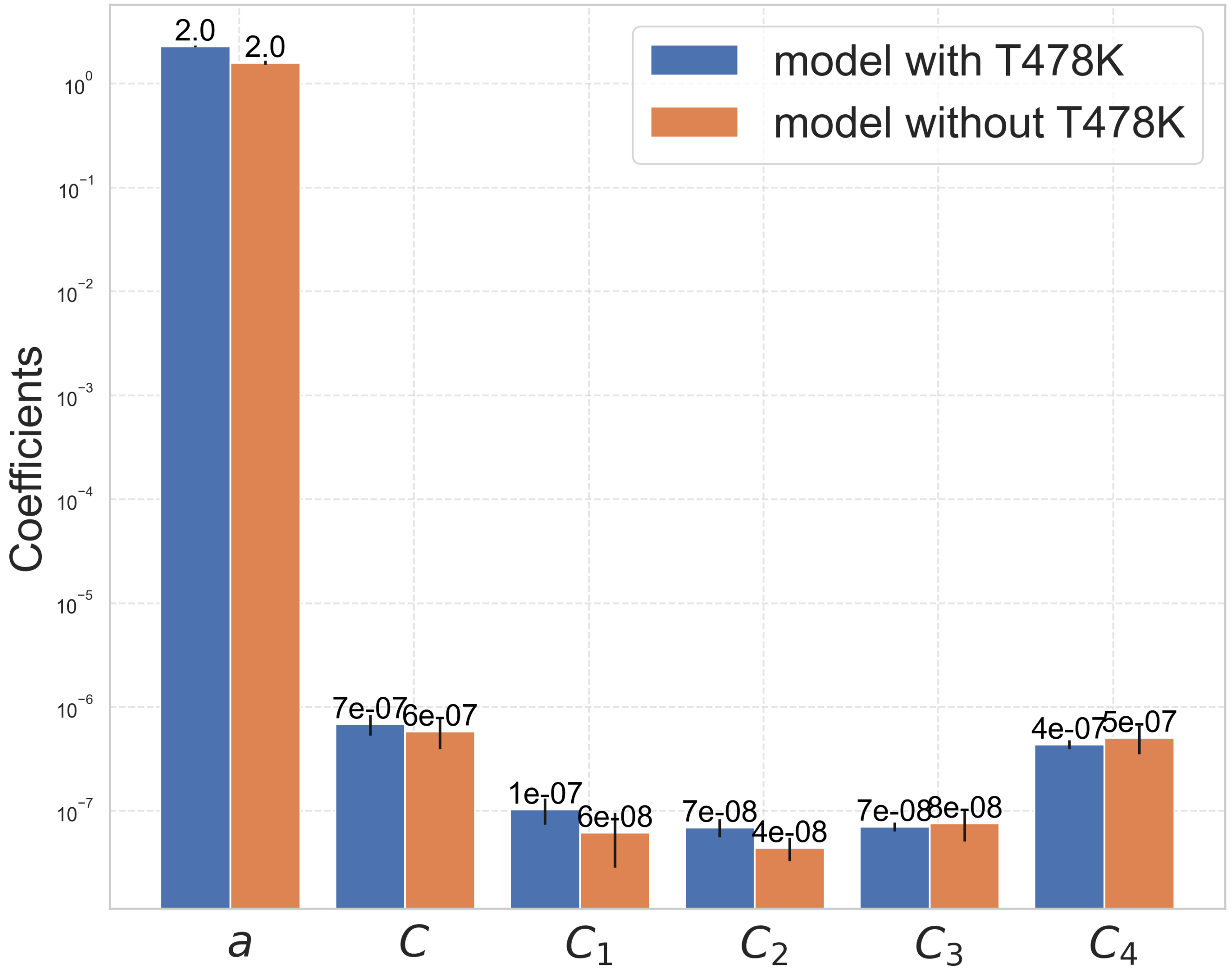


Fig. S.4. Robustness of the model: bar plot of the fitted coefficients from the biophysical model, presenting variants with and without the T478K mutation.

##### S4. *Energy Scale as a hyperparameter*

We selected a value of $T = 1.6$ to ensure that the fitted effective concentrations align with the values of antibody concentrations observed in nature [16]. The fitted parameters as a function of temperature are shown in Figure S.5a.

It is crucial to note that the choice of $T$ exerts minimal influence on the model's behavior, as demonstrated in Figure S.5b-f. The model behaviors remains stable as we alters $T$, and it is even less susceptible to change in the regions where nature has explored the $K_d$ values.

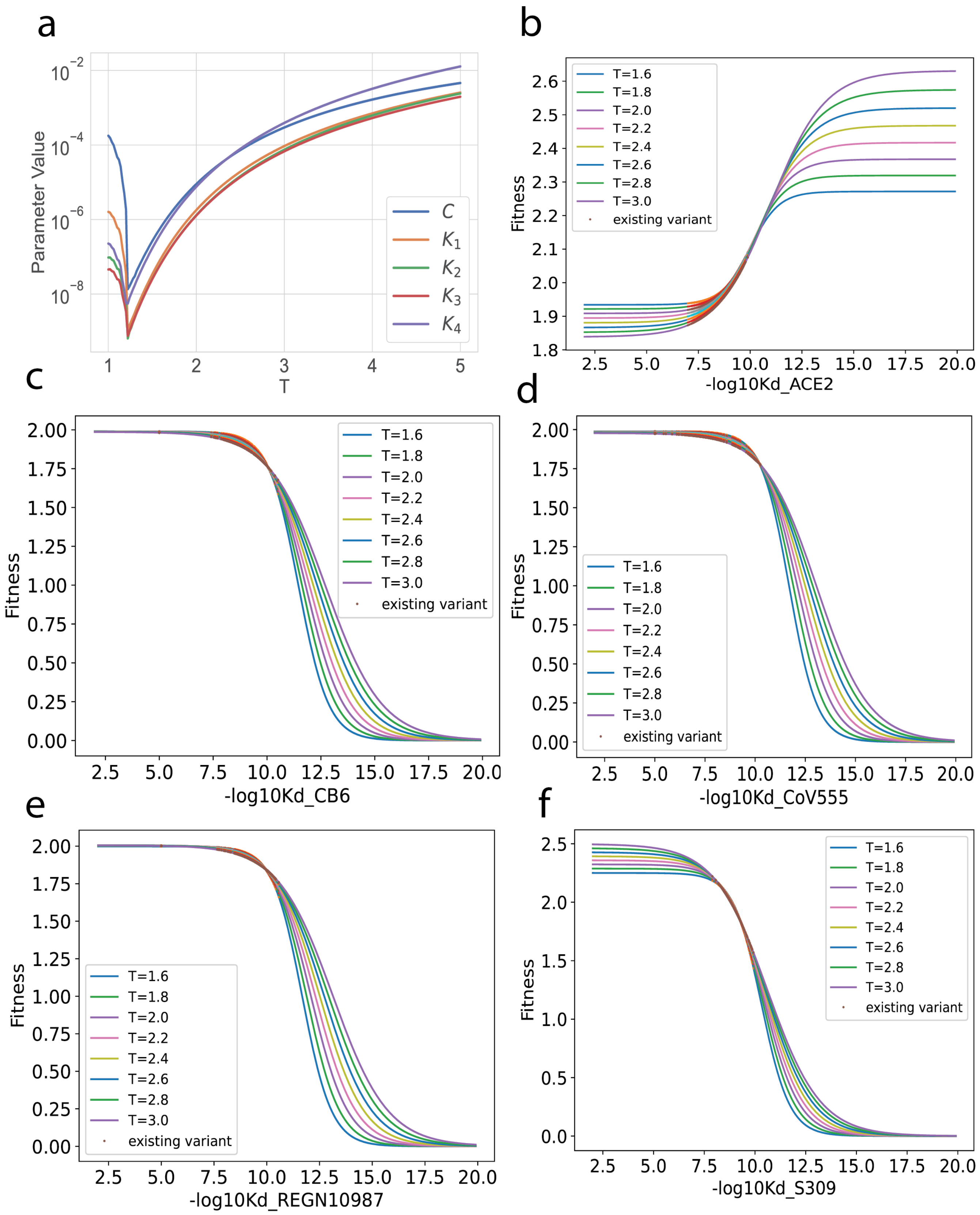


Fig. S.5. Effect of Energy Scale hyperparameter. In (a): Parameters obtained from the logistic regression as a function of T. In (b-f): Relationship between fitness and binding constants at different Energy Scales. The fitted curve is slightly affected by changes in T, on domains where existing variants fall (red dots).

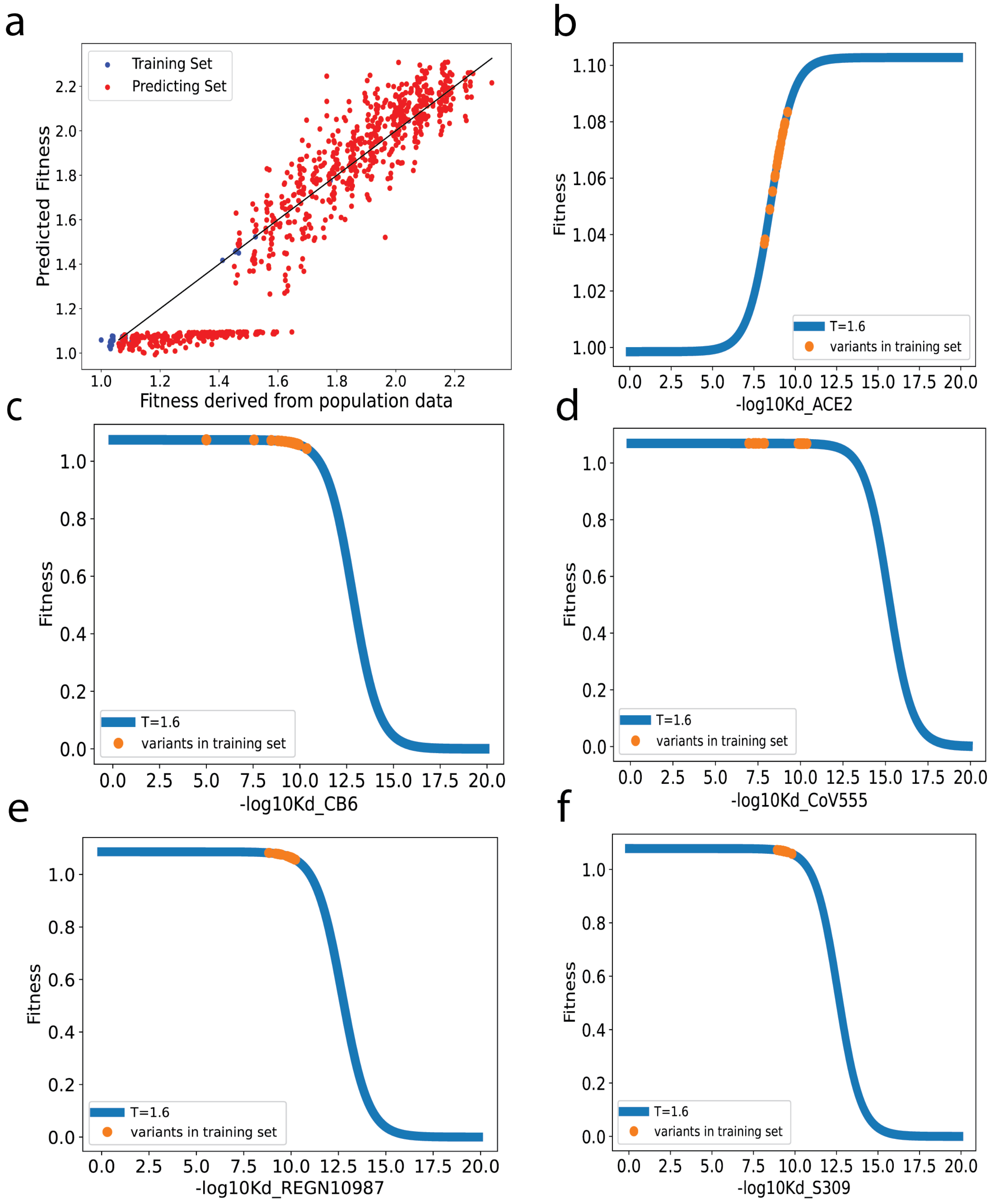


Fig. S.6. Prediction of future variants: (a) The comparison between predicted fitness values and true fitness values, with the training set cut-off in May 2021. The testing set contains all the variants between May 2021 and May 2022 (b-f) The relationship between fitness and binding constants as fitted by our model, for variants that appeared before May 2021 and that lack the T478K mutation.

In Fig S.6a, we trained our model using 8 data points with the T478K mutation and 34 points without it. Impressively, we were able to accurately predict variants carrying the T478K mutation using only these 8 training points, contributing to our successful prediction of fitness jump caused by the Omicron BA.1. variant.

However, a systematic underestimation was observed for variants with lower fitness, for those lacking the T478K mutation. This underestimation can be ascribed to the fact that variants without the T478K mutation in the training set failed to exhibit a fitness advantage even when escaping antibodies. Note that this characteristic is demonstrated in Fig S.6c-f, where all the data points reside within the upper plateau region of the logistic curve.

#### *S6. Justification of the split of the data using T478K and biophysical implications*

Our proposed model posits that fitness can be depicted as a biophysical function of disassociation constants to cell receptors and antibodies, assuming the virus's fitness is primarily influenced by its ability to bind to the cell receptor (equation (1)). Our initial attempt to fit a biophysical model to the entire dataset, encompassing the dissociation constants and fitness of observed variants, shows data can be divided into two subsets, each of which following a unique trend (Figure S.7). An in-depth analysis of these subsets indicates that they are primarily distinguished by the presence or absence of the mutation T478K.

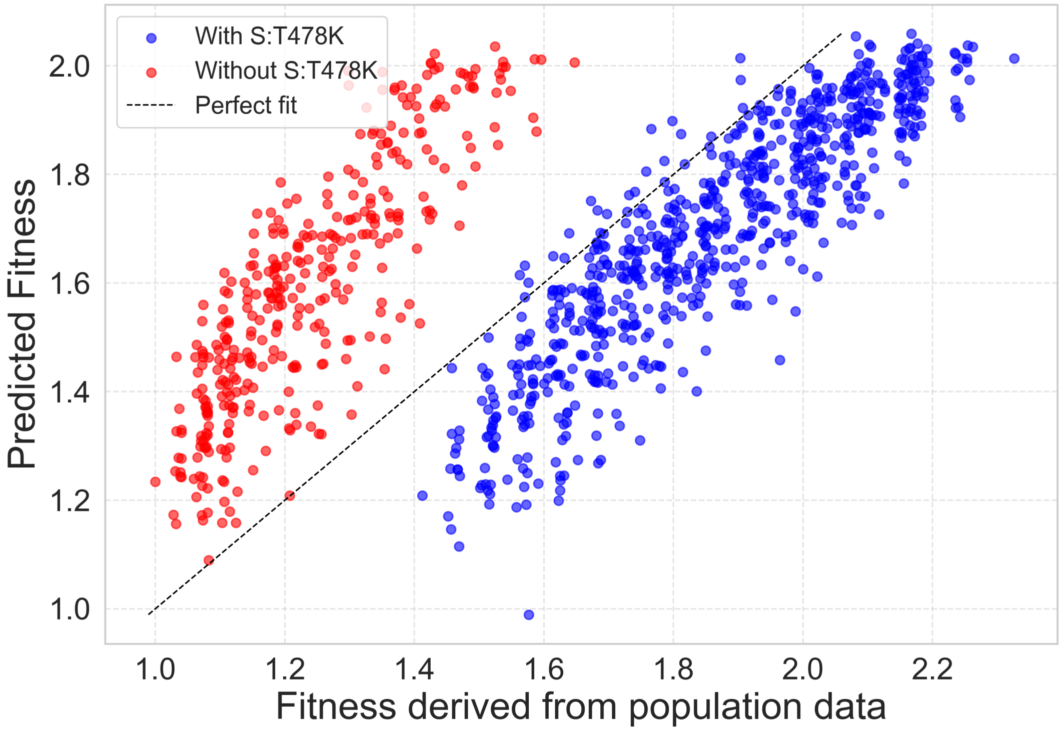


Fig. S.7. Comparison of predicted fitness vs true fitness for existing variants with and without Mutation T478K: two distinct trends can be observed.

In this section, we motivate why the data were split based on the presence of the T478K mutation and why we observe different scaling parameters on each subset. Our model implies fitness $F$ can be written as a biophysical function $\Psi$ of the biophysical parameters $k_i, i \in [1,5]$, times a scaling factor $a$, for each dataset where mutations fitness increase is due to improvement of biophysical parameters: $F = a\Psi(k_i)$

For each variant with mutations $X_m$ and log of binding constants $k_i$, this can be written for the dataset without T478K :

$$\exp\left(\sum_{m,m\in RBD,m\neq T478K} b_m X_m\right) = a\Psi(k_i)$$

For each variant with mutations $X'_m$ and binding constants $k'_i$, this can be written for the dataset with T478K :

$$\exp\left(b_{T478K} + \sum_{m,m\in RBD,m\neq T478K} b_m X'_m\right) = a'\Psi'(k'_i)$$

In addition, following the study of Moulana et al. [8,9] that showed T478K did not impact binding constants, even by co-occurence with other mutations, we get that adding or removing T478K mutation (while keeping the other mutations $X_m$) does not change the binding constants $k_i$. Formally: $\forall m \neq T478K, X_m = X'_m \implies \forall i, k_i = k'_i$

Now let's consider an undetermined variant with mutations $X_m$ outside T478K. Noting that $\exp(b_{T478K}+\sum_{m,m\in RBD,m\neq T478K} b_m X_m) = e^{b_{T478K}} \exp(\sum_{m,m\in RBD,m\neq T478K} b_m X_m)$, our model implies:

$$a'\Psi'(k'_i) = e^{b_{T478K}} a\Psi(k_i)$$

Since $k_i = k'_i$, we can expect:

$$\Psi = \Psi' \text{ and } a' = e^{b_{T478K}} a$$

In other words, presence of T478K should only impact the scaling factor of the biophysical model. Reminding that $e^{b_{T478K}} \approx 1.41$ [15], we check that the ratio between scaling factors in Figure 2d is $2.27/1.58 \approx 1.43$, which is close to 1.41, and that fitted concentrations are roughly the same.

Thus, splitting data on the presence of mutation T478K enables to incorporate the impact of T478K on fitness in the scaling factor, but doesn't change the relationship between binding constants and fitness: the mapping between biophysical parameters and fitness that is inferred by our model remains the same.
